# Dopaminergic D2 receptor modulation of striatal cholinergic interneurons contributes to sequence learning

**DOI:** 10.1101/2023.08.28.554807

**Authors:** Jessica Hotard Chancey, Christoph Kellendonk, Jonathan A. Javitch, David M. Lovinger

**Author notes:** Address correspondence to: David Lovinger, National Institutes on Alcohol Abuse and Alcoholism National Institutes of Health, 5625 Fishers Lane, Room TS-13A Rockville, MD 20852.

## Abstract

Learning action sequences is necessary for normal daily activities. Medium spiny neurons (MSNs) in the dorsal striatum (dStr) encode action sequences through changes in firing at the start and/or stop of action sequences or sustained changes in firing throughout the sequence. Acetylcholine (ACh), released from cholinergic interneurons (ChIs), regulates striatal function by modulating MSN and interneuron excitability, dopamine and glutamate release, and synaptic plasticity. Cholinergic neurons in dStr pause their tonic firing during the performance of learned action sequences. Activation of dopamine type-2 receptors (D2Rs) on ChIs is one mechanism of ChI pausing. In this study we show that deleting D2Rs from ChIs by crossing D2-floxed with ChAT-Cre mice (D2Flox-ChATCre), which inhibits dopamine-mediated ChI pausing and leads to deficits in an operant action sequence task and lower breakpoints in a progressive ratio task. These data suggest that D2Flox-ChATCre mice have reduced motivation to work for sucrose reward, but show no generalized motor skill deficits. D2Flox-ChATCre mice perform similarly to controls in a simple reversal learning task, indicating normal behavioral flexibility, a cognitive function associated with ChIs. *In vivo* electrophysiological recordings show that D2Flox-ChatCre mice have deficits in sequence encoding, with fewer dStr MSNs encoding entire action sequences compared to controls. Thus, ChI D2R deletion appears to impair a neural substrate of action chunking. Virally replacing D2Rs in dStr ChIs in adult mice improves action sequence learning, but not the lower breakpoints, further suggesting that D2Rs on ChIs in the dStr are critical for sequence learning, but not for driving the motivational aspects of the task.

**Significance statement:** The role of striatal projection neurons in encoding action sequences has been extensively studied, and cholinergic interneurons play a central role in striatal physiology, but we do not yet understand how cholinergic interneurons contribute to action sequencing. Using a combination of mouse genetics, behavior, and *in vivo* electrophysiology this work shows that genetic deletion of D2 receptors from striatal cholinergic interneurons disrupts the learning, performance, and encoding of action sequences, without changing general locomotion or motor skill learning. Virally replacing D2 receptors specifically in dorsal striatal cholinergic interneurons is sufficient to rescue the sequence behavior. Our observations may be useful in understanding and treating movement disorders in which dopamine and acetylcholine are imbalanced.

## Introduction

Learning action sequences is necessary for normal daily activities, such as tying your shoes. With extended training individual movements are chunked together, such that an individual can complete an entire sequence without consciously thinking about the separate actions necessary to complete the sequence (*1*). The dorsal striatum (dStr) plays a critical role in action selection, action initiation, and encoding of learned action sequences. Specifically, dStr medium spiny neurons (MSNs) change their firing rate at the start and/or termination of action sequences (*2–5*). MSNs also encode entire learned action sequences as a single event through sustained changes in firing throughout the sequence in well trained animals (*3*).

The striatal network is modulated by the actions of acetylcholine (ACh), which is released from tonically active cholinergic interneurons (ChIs). ChIs are only a small subset (1-2%) of the total neurons in the striatum, but their dense axonal branches release ACh that acts on a diverse group of muscarinic and nicotinic ACh receptors found on most cells and terminals throughout the striatum. This cholinergic transmission modulates the firing of MSNs and GABAergic interneurons, and neurotransmitter release from glutamatergic and dopaminergic terminals (*6, 7*). Thus, ACh is important for many striatal-dependent behaviors and synaptic plasticity. ChIs also corelease glutamate (*8, 9*) and are themselves heavily regulated. They receive glutamatergic input from the cortex and thalamus, GABAergic input from MSNs, interneurons and extrastriatal inputs, ACh from neighboring ChIs, and dopamine from the midbrain (*7, 10, 11*). An imbalance between the cholinergic and dopaminergic systems is thought to contribute to the symptoms of movement disorders such as Parkinson’s and Huntington’s Diseases (*12, 13*).

The physiology of ChIs is regulated by dopamine, including via dopamine type-2 receptors (D2Rs) that decrease tonic firing and reduce ACh release (*14, 15*).The D2Rs expressed on ChIs also participate in long-term synaptic depression thought to be involved in striatal based learning (*16*). There are very few *in vivo* recordings from ChIs in mice, but Jin et al. (2014) found that ChIs in the dStr decrease, or pause, their tonic firing during the performance of a learned sequence of lever pressing (3). ChIs are thought to be analogous to tonically active neurons (TANs) in primates (*17*), which have been shown to pause their firing in response to learned motor responses and conditioned stimuli (*18, 19*). Similarly, stimulus-induced pauses in firing have been reported in mouse ChIs (*20*). This pausing physiology has been attributed to D2R activation *in vivo* in primates (*21*), and activation of D2Rs on ChIs drives pauses in firing in brain slices from mice (*14, 22–25*). Furthermore, D2Rs contribute to decreases in ACh levels induced by environmental stimuli and during performance of a reward-based decision making task in vivo (28, 29, 30). However, other mechanisms can induce pauses in ChI firing (28) (*26*). Loss of D2Rs in ChIs has been shown to reduce the locomotor response to cocaine (29) (*27*), but the significance of this dopamine-mediated regulation of ChIs for self-directed behavioral performance has not been explored.

We hypothesized that D2R modulation of ChIs, which produces pausing or alterations in synaptic plasticity, is critical to regulate MSN activity necessary for action and sequence learning. We found that genetic deletion of D2Rs from ChIs leads to deficits in operant lever pressing and progressive ratio tasks, without changing rotarod performance or behavioral flexibility. We also found that loss of D2Rs in ChIs produces deficits in sequence encoding, with fewer MSNs encoding entire action sequences compared to controls, suggesting that ChI D2R deletion impairs a neural substrate of action chunking. We show that virally replacing D2Rs in ChIs in the dStr, improves action sequence learning, but not the lower breakpoints, further suggesting that D2Rs on ChIs in the dorsal striatum are critical for sequence learning, but not for driving the motivational aspects of the task.

## Results

To study dopaminergic modulation of ChIs, we genetically deleted D2Rs from ChIs by crossing D2R-floxed (D2flox) mice with mice that express the Cre recombinase under the control of the choline acetyltransferase promoter (ChATCre) to make D2flox-ChATCre mice. We have shown that this leads to D2R loss specifically on ChIs within the striatum and that dopamine-mediated pausing of ChI firing is lost in these mice, measured in *ex vivo* slice recordings (*24*). Genetic deletion of D2Rs from ChIs also reduces cue-evoked dips in acetylcholine levels *in vivo* (26) (*28*), and overexpression of D2Rs in ChIs increases dopamine mediated pause duration (27) (*29*).

Because the dorsal striatum is important for the learning and execution of action sequences, we tested D2flox-ChAT-IRES-Cre mice in a self-paced lever-pressing action sequence paradigm adapted from Jin and Costa (*2*). In this paradigm, mice were required to press a lever 8 times to receive a single sucrose reinforcer, but the count was re-set when animals attempted to retrieve the sucrose prior to completing the pressing sequence. The training progressed from fixed-ratio (FR)1 to FR4 to FR8 for a minimum number of training days (3, 4, and 8 days, respectively) or until they reached a criterion (see methods for full description). This resulted in rapid learning of sequences with low variability in wild-type C57/B6 mice (Fig. S1). D2flox-ChATCre animals, however, did not perform as well as D2flox littermate controls in the sequence task. Fig. 1A shows examples of pressing behavior in control and knockout animals, in which the D2flox animal pressed in consistent sequences with a high rate of reward attainment on the fourth day of FR4 (Fig. S2) and the eighth day of FR8 training (Fig. 1A, left). In contrast, the D2flox-ChATCre littermate pressed fewer times and with lower efficiency ([#rewards earned/sequences] *100) (Fig. 1A, right). Summary data are shown in Fig. 1B-G. D2flox-ChATCre mice required more training to reach criteria (15 rewards in a 1 hour session) throughout the paradigm (Fig. 1B), but eventually reached the same level of efficiency as controls with additional training (Fig. 1C). Similarly, they pressed fewer times per sequence, but reached levels similar to controls with increased training (Fig. 1D). Although they reached a level of efficiency similar to controls, D2flox-ChATCre animals received fewer rewards throughout training (Fig. 1D), initiated fewer sequences, as demonstrated by the increased inter-sequence interval (Fig. 1E) and pressed at a slower rate within each pressing sequence (Fig. 1G) than D2flox littermates.

**Figure 1.**
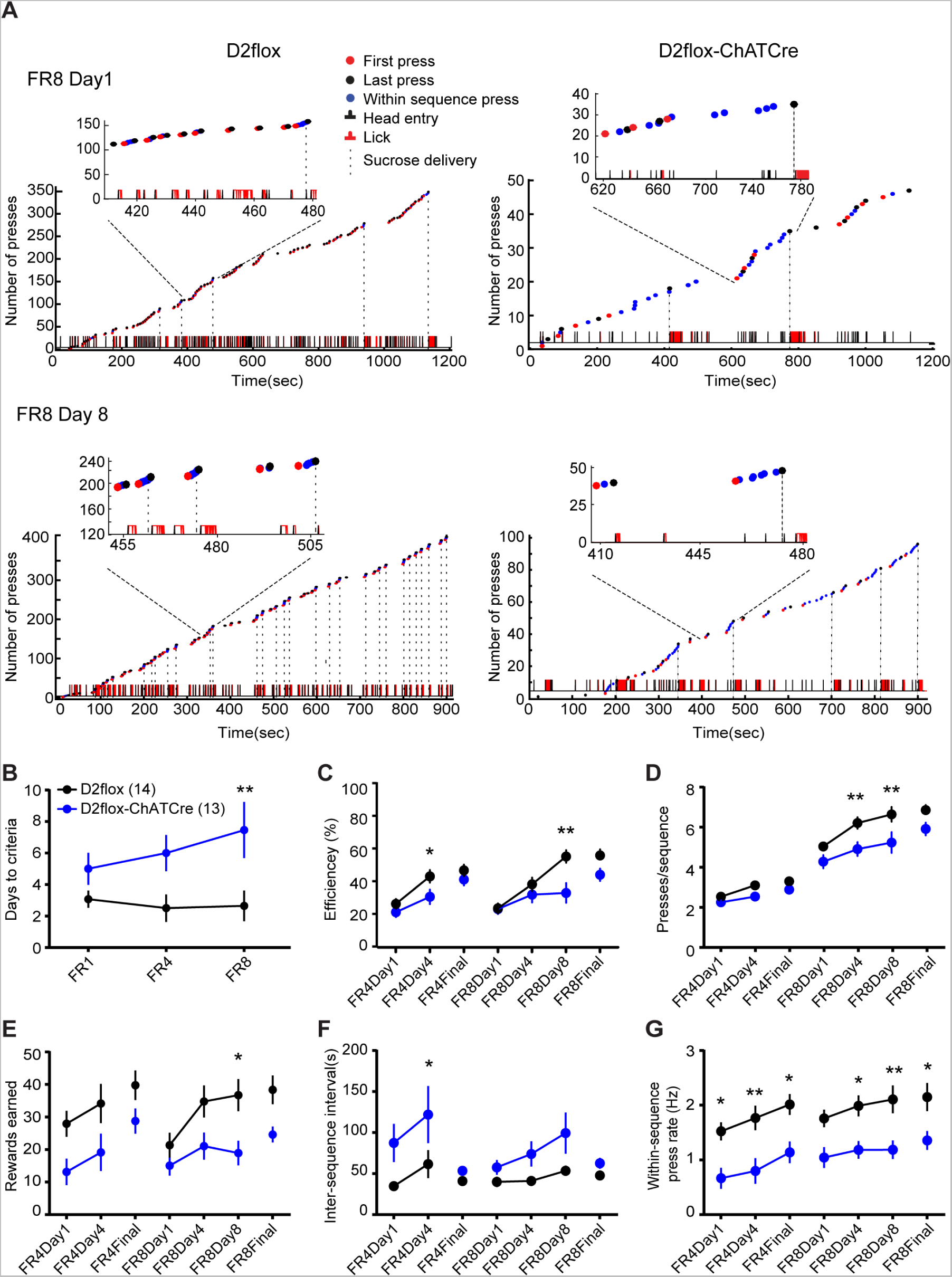
D2flox-ChATCre animals have deficits in sequence learning. **A)** Cumulative lever press plots showing examples of behavioral shaping in D2flox (left) and D2flox-ChATCre (right) littermates performing the adapted FR task. Top: The first day of FR8 training. Bottom: 8^th^ day of FR8 training in which this D2flox-ChATCre animal presses less frequently, slower, and with lower efficiency than the D2flox littermate. **B)** D2flox-ChATCre (blue) mice required more training than littermates (black) as measured by days to reach the criterion (main effect of genotype: F_(1,25)_ = 16.6, p < 0.001, two-way repeated measures ANOVA). *A priori* Sidak’s multiple comparisons tests between genotypes at each stage of training were performed in B-G. Significant differences are indicated by an asterisk (* p < 0.05, ** p < 0.01). **C)** D2flox-ChATCre animals reached similar levels of efficiency (% of sequences that are rewarded) as control littermates but did so at a slower rate (main effect of training day F_(6,150)_ = 18.97, p < 0.01; genotype: F_(1,25)_ = 7.41, p < 0.05; interaction F_(6,150)_ = 2.29, p < 0.05). **D)** Presses per sequence increased with training (main effect of training day: F_(6,150)_ = 109.60, p < 0.0001) in both genotypes, but were lower in D2flox-ChatCre (main effect of genotype: : F_(1,25)_ = 10.49, p < 0.01). **E)** Number of reinforcers increased with training (main effect of training day: F_(6,150)_ = 5.86, p < 0.0001) in both genotypes, but D2flox-ChatCre received fewer reinforcers per session throughout training (main effect of genotype: : F_(1,25)_ = 8.78, p < 0.01). **F)** The inter-sequence interval, or time between onset of pressing sequences, was longer in D2flox-ChATCre animals compared to controls (main effect of genotype: F_(1,25)_ = 7.85, p < 0.01). **G)** The within-sequence press rate also increased throughout training (main effect of training day: F_(6,150)_ = 8.26, p < 0.0001) in both genotypes, but was lower in D2flox-ChatCre (main effect of genotype: : F_(1,25)_ = 12.26, p < 0.01).

There are two ChATCre mouse lines, and both have been shown to have minor, but differential behavioral deficits (*31*). To confirm that the behavioral deficits reported thus far are due to loss of D2Rs from ChIs, we trained a second D2flox-ChATCre line formed by crossing D2flox with Tg-BAC-ChATCre mice. These mice showed similar deficits in sequence learning as the D2flox-Chat-IRES-Cre mice (Fig. S3), including lower efficiency, presses per sequence, rates of pressing, and rewards earned, and required additional training to reach criteria compared to littermate controls. The *ChAT* locus also contains the gene for the vesicular acetylcholine transporter (VAChT) (*32*), and another BAC transgenic ChAT mouse line that expresses channelrhodopsin under the control of the ChAT promoter (ChAT-ChR2(H134R)-EYFP) have been shown to have excessive VAChT expression and motor abnormalities (*33, 34*). We, therefore, also examined the performance of Tg-BAC-ChATCre mice (hemizygotes compared to Cre-littermates) in the sequence paradigm. We found that Tg-BAC-ChATCre^+^ animals performed as well as their Cre negative littermates in the sequence pressing paradigm (Fig. S4A-D). Additionally, the Tg-BAC-ChATCre animals performed similar to wild-types on an accelerating rotarod task (Fig. S4E), a paradigm that was disrupted in ChAT-ChR2(H134R)-EYFP animals (*33*), indicating that the sequence learning deficits in the D2flox-BAC-ChATCre animals were due to D2R deletion from ChIs and not aberrant VChAT expression.

Our findings, with two independent D2flox-ChATCre lines, show that removing ChI D2Rs caused a deficit in performing self-paced action sequences. Knockout animals were able to learn the action sequences and performed with similar efficiency to controls with additional training. A second phenotype present in both lines that did not normalize to control levels with increased training was a decrease in the amount of pressing, leading to fewer earned rewards. There are several possible reasons for these phenotypes including motor deficits, motivational deficits, or deficits in behavioral flexibility, an aspect of cognitive function impaired in ChI ablation studies. We therefore examined these possibilities in D2flox-ChATCre mice. In later experiments, unless otherwise stated, we combined data from the two D2flox-ChATCre lines, after confirming there was no significant difference between the two lines. To test for gross motor deficits, we placed the animals in a novel cage and monitored locomotor activity using beam breaks. As expected, we saw novelty-induced locomotor activity early in the session, and locomotion decreased throughout the session (Fig. 2A). There was no difference between genotypes, indicating that D2flox-ChATCre mice do not have gross motor deficits. Likewise, D2flox-ChATCre animals performed similar to D2flox littermates in an accelerating rotarod task (Fig. 2B), indicating that they do not have generalized deficits in motor skill learning.

**Figure 2.**
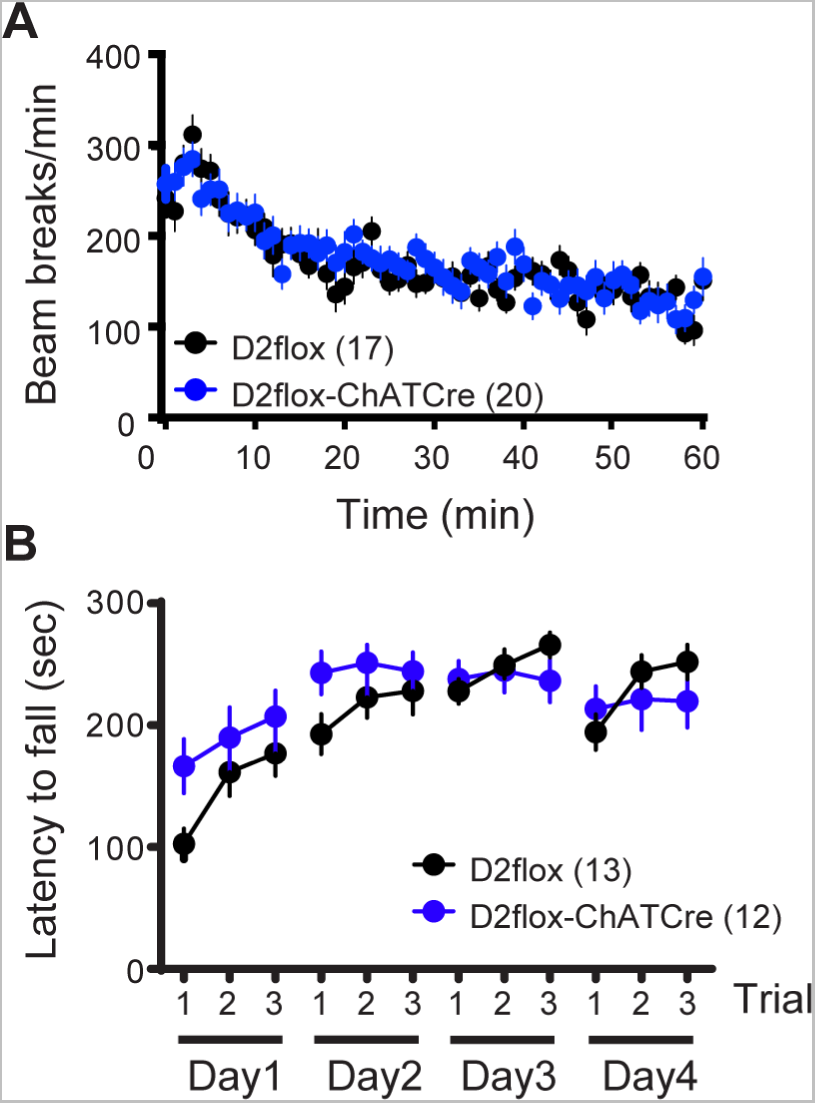
D2flox-ChATCre mice do not have motor deficits. **A)** Mice were placed in a novel cage, and locomotor activity was measured via beam breaks. Both genotypes showed heightened novelty induced locomotion, and a decrease in motion as the one hour session progressed (main effect of time: F_(60,2100)_ = 18.33, p < 0.001, two-way repeated measures ANOVA). Activity was similar between genotypes (F_(1,35)_ = 0.1197, p = 0.73). N values in parentheses. **B)** D2flox-ChATCre and D2flox littermates learned the accelerating rotarod task with a similar time course and performance level (main effect of time: F_(11,253)_ = 9.40, p < 0.001; genotype : F_(1,23)_ = 0.62, p = 0.44).

We next tested reversal learning, because ablating ChIs (*35, 36*), or removing the thalamic inputs to ChIs (*37*) has been shown to interfere with behavioral flexibility, and the sequence task required flexible learning. We ran a simple self-paced lever reversal paradigm in which animals learned to press on two different reinforced levers. After an initial training phase in which both levers were reinforced, a single lever was reinforced, while the other was not (acquisition phase, Fig. 3A-B), and midway through training, the reinforced lever was reversed. Reinforcement was delivered on an FR1 schedule to keep the effort low, because D2flox-ChATCre animals appeared to decrease their performance with increased work levels in the sequence task (Fig. 1). We found that D2flox-ChATCre animals performed similar to controls in this simple reversal task. There was no difference between genotypes in the proportion of reinforced presses (Fig. 3A) or in perseverative errors (not shown). As in the sequence task, D2flox-ChATCre mice appeared to receive fewer rewards than controls (Fig. 3B), but this difference was not significant. Because we repeatedly saw that D2flox-ChATCre animals received fewer rewards than controls (Figs. 2E, S4C, 4B), we tested the rewarding properties of sucrose using a home cage sucrose preference test, in which naïve mice were given 10% sucrose and water, and we measured the ratio of sucrose consumed compared to water in a 6 hour period. We found that D2flox-ChATCre mice exhibited preference similar to control littermates (Fig. 3C), indicating that the reduced number of rewards earned in the operant tasks was not due to a reduced hedonic value of sucrose. We therefore tested the motivation to work for sucrose using a self-paced progressive ratio operant paradigm. D2flox-ChAT-IRES-Cre mice pressed fewer times than D2flox controls (Fig. 3D) and had lower breakpoints (Fig. 3E). A similar phenotype was found in D2flox-BAC-ChATCre mice (Fig. S3 G-I). Together these data indicate that mice lacking D2Rs in ChIs display a reduced motivation to work for sucrose reward.

**Figure 3.**
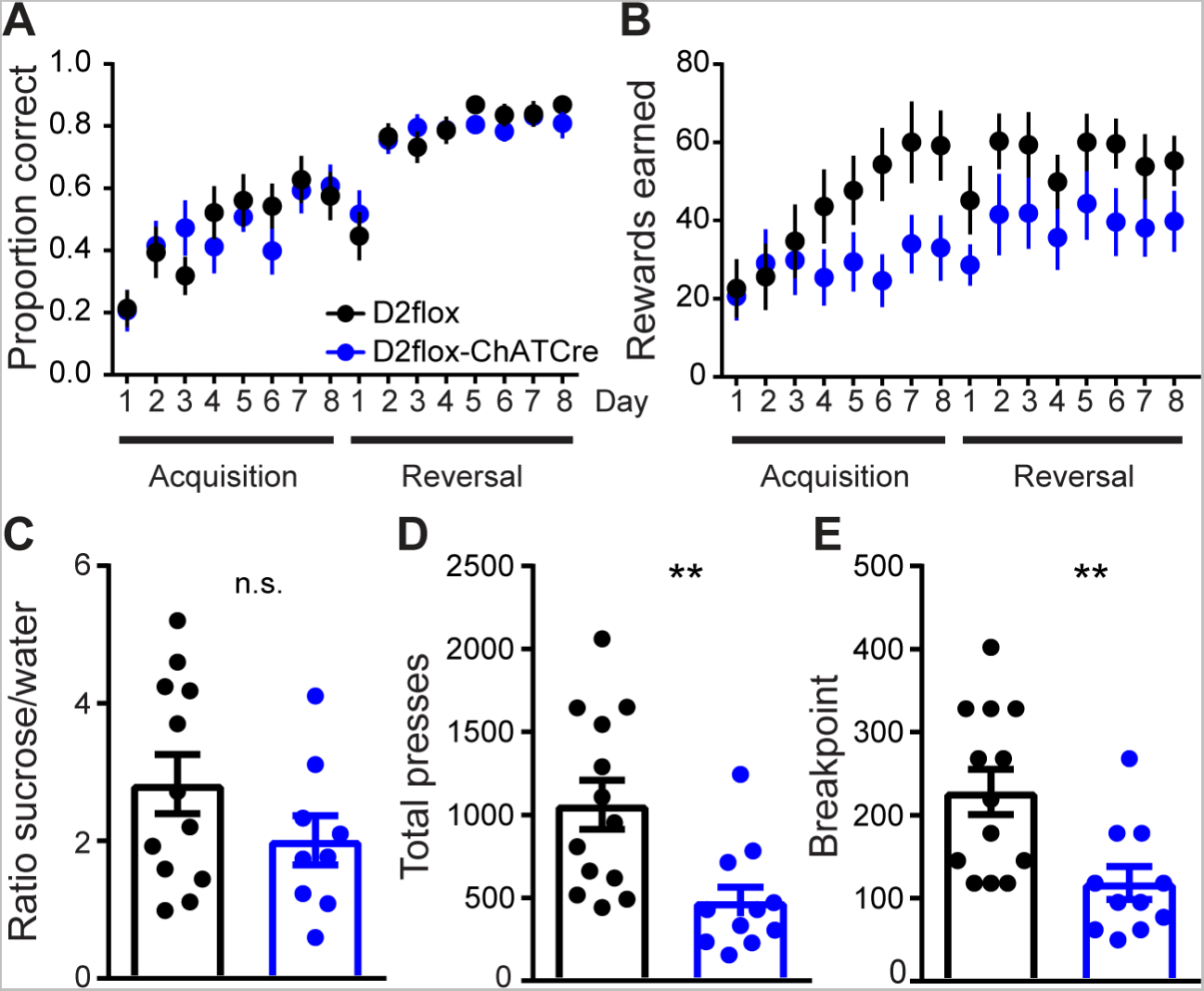
D2flox-ChATCre mice perform normally in a flexibility task but have lower breakpoints. **A)** Lever pressing in the discrimination and reversal phases of a self-paced two lever instrumental learning task in D2flox (black; n = 11) and D2flox-ChATCre mice (blue; n = 11) were similar. Both groups learned the reversal task, as evidenced by an increase in the proportion of rewarded lever presses (main effect of training day: F_(15,300)_ = 22.77, p < 0.001; no effect of genotype: F_(1,20)_ = 1.30, p = 0.72). **B)** No significant difference between the two groups was observed in rewards received throughout the task (main effect of training day: F_(15,300)_ = 6.10, p < 0.001; genotype: F_(1,20)_ = 3.05, p = 0.09). **C)** In a home cage sucrose preference test, both genotypes showed a similar preference for sucrose (D2flox: 2.83 ± 0.43, n = 12; D2flox-ChATCre: 2.01 ± 0.36, n = 9; p < 0.01 compared to 1 [i.e., no preference] for both genotypes; p = 0.18 comparing genotypes, Student’s t-test). **D)** In a progressive ratio paradigm, D2flox-ChATCre mice had lower rates of total pressing (481.5 ± 96.3; n = 13) compared to D2flox littermates (1061.0 ± 147.8; p < 0.01; n = 11), as well as lower breakpoints (**E)** D2flox: 227.9 ± 27.3; D2flox-ChATCre: 118.3 ± 19.9; p < 0.01).

**Figure 4.**
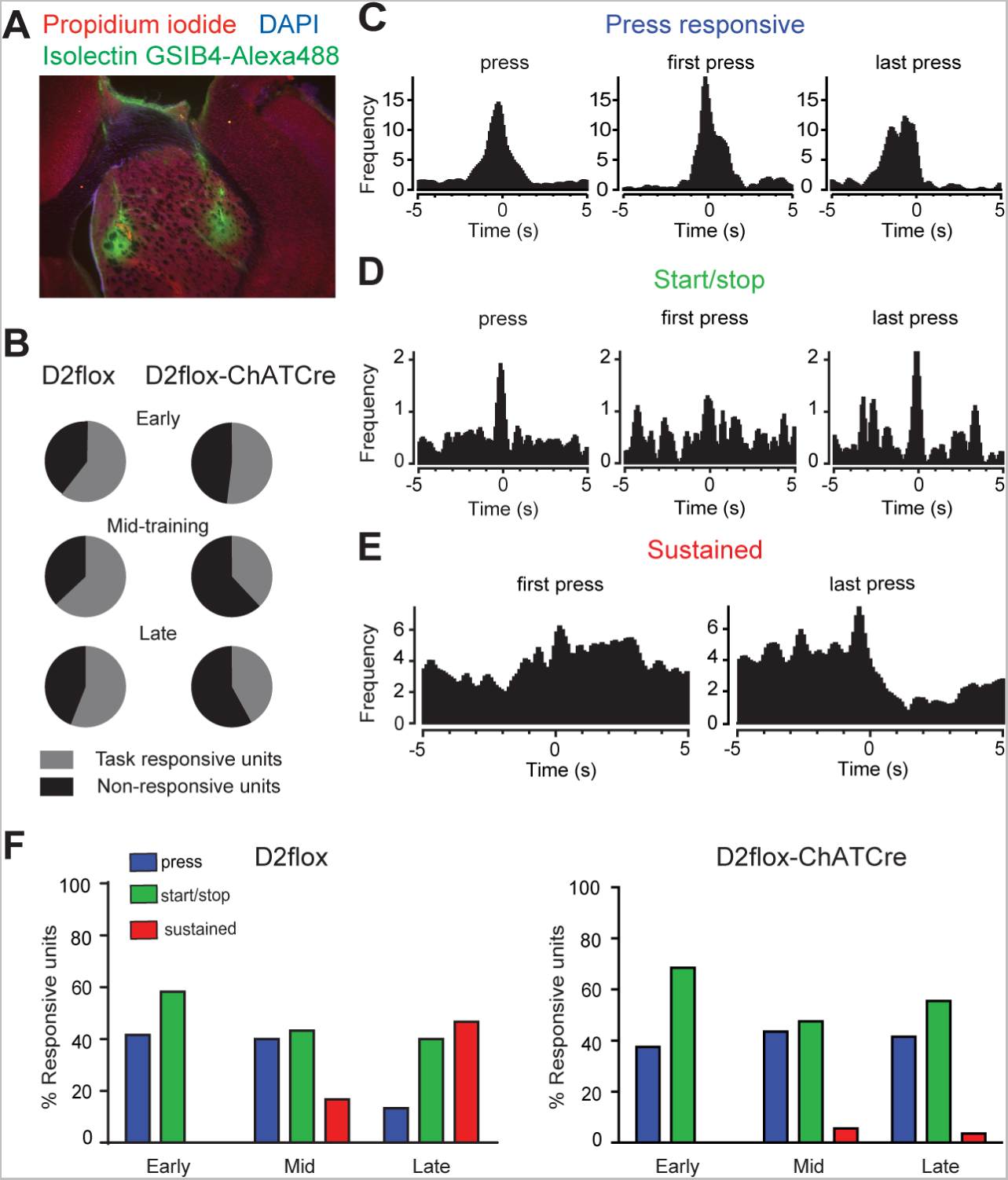
Sequence encoding by MSNs is disrupted in D2flox-ChATCre mice. **A)** 16 channel micro-arrays were implanted into dStr (8 in DMS and 8 in DLS). Electrode placement was verified by tract staining with Isolectin GS-IB4. Scale = 220 μm. **B)** Similar percentage of task responsive putative MSNs across genotypes (n = 84 units from 7 animals D2flox; n = 71 units from 7 animals D2flox-ChATCre; p > 0.05; χ2). **C)** Examples of press responsive (top), **D)** start/stop (middle) and **E)** sustained chunking (bottom) putative MSNs. **F)** D2flox-ChATCre animals have lower percentage (1 of 21 units) of task responsive “chunking” units that have sustained changes in firing throughout the sequence of pressing compared to D2flox littermates (11 of 25 units; p < 0.01; χ2).

To better understand the deficit in action sequencing we examined how removing D2Rs from ChIs affects neuronal encoding of action sequences. Jin and colleagues (*2, 3*) found that MSNs in the dorsal striatum encode the start, stop, and concatenation of action sequences. We repeated our sequence paradigm in D2flox and D2flox-ChATCre mice implanted with recording electrodes in the dorsal medial and dorsal lateral striatum (Fig. 4A). Similar to the previous results, D2flox-ChATCre mice did not perform as well as D2flox littermates (Fig. S5), and all implanted animals were slower to learn than nonimplanted animals. As previously shown, a substantial proportion of neurons showed activity that correlated with some aspect of the behavior (task responsive units, Fig. 4B). With increased FR8 sequence training, the percentage of task responsive units did not change significantly (Fig. 4B). From the beginning of FR8 sequence training MSNs in the DS showed altered firing in relation to single lever presses withing sequences (Fig. 4C), or at the start and stop of pressing sequences (Figs. 4D and S6). As training progressed we observed the appearance of “action chunking” cells, or units that change their firing rate throughout the performance of the entire action sequence (Fig. 4E-F). These chunking cells were only found in the lateral portion of the dorsal striatum. Interestingly, D2flox-ChATCre animals displayed similar encoding of action onset and termination, but units with sustained action chunking activity were nearly absent in the knockout animals, even with additional training and a similar efficiency of performance (Fig. 4F). These data indicate that removal of D2 receptors from ChIs impairs the ability of MSNs to encode the full sequence duration, which could contribute to poor sequence performance.

To determine if the behavioral deficits were indeed due to the absence of D2 receptors in dorsal striatal ChIs rather than any developmental abnormalities, we virally replaced D2 receptors in dorsal striatal ChIs of adult mice using a Cre-dependent viral construct carrying the *drd2* gene (*37*). AAV2/1-hSyn-DIO-D2R(L)-IRES-mVenus or AAV2-DIO-eGFP was injected into the dStr of D2flox-ChATCre mice or D2flox littermates (Fig. 5A), and viral expression was limited to ChIs (Fig. 5B). As shown previously, D2flox-ChATCre mice expressing the eGFP control virus (KO-GFP, Fig. 5) had deficits in performance of the sequence and progressive ratio tasks compared to D2flox littermates injected with the control GFP virus (Con-GFP). The behavior of D2flox-ChATCre injected with the D2 virus normalized to that of controls in the sequence task, indicating that D2 receptors in dStr ChIs are important for the learning and performance of this behavior. However, the viral rescue animals still showed a deficit in the progressive ratio task, indicating that the loss of D2Rs in dStr ChIs is likely not driving this behavior or that that adult rescue is unable to overcome a developmental phenotype.

**Figure 5.**
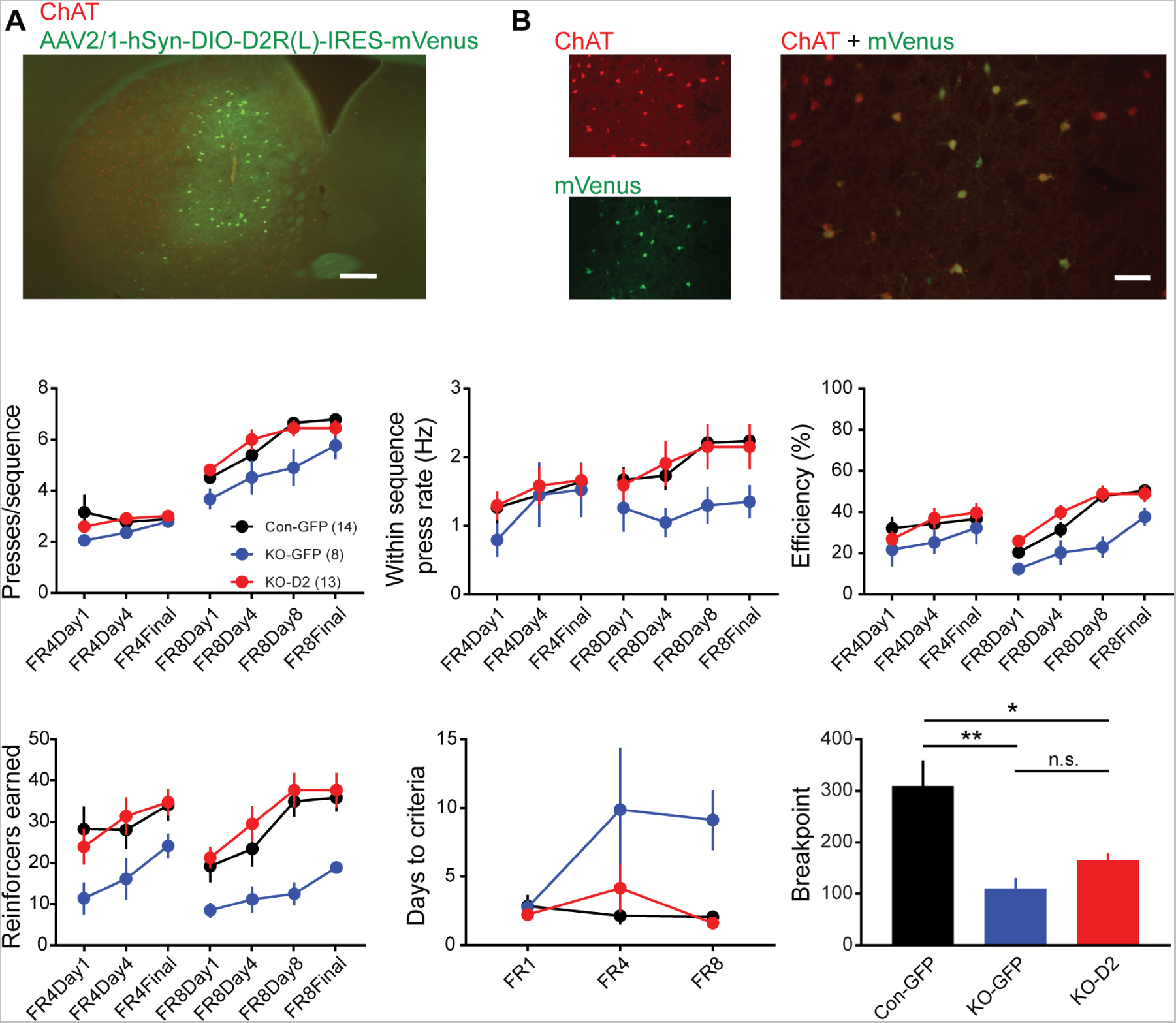
Virally replacing D2Rs in dorsal striatal ChIs rescues sequence learning but not the progressive ratio task. **A)** Example of AAV2/1-hSyn-DIO-D2R-IRES-mVenus virus (green) injected into the dorsal striatum of a D2flox-ChATCre mouse that was stained for choline transferase (ChAT, red). Scale = 220 μm. **B)** Viral expression was limited to ChAT+ cells. Scale = 70 μm. **C-G)** Con-GFP (black) are D2flox animals injected with the control virus (AAV2-DIO-eGFP). KO-GFP (blue) are D2flox-ChATCre mice injected with AAV2-DIO-eGFP. KO-D2 (red) are the rescue group in which D2flox-ChATCre mice were injected with the D2 rescue virus. N values in parentheses. Two-way repeated measures ANOVAs were performed with posthoc Dunnett’s multiple comparisons tests comparing the KO groups to the control group (Con-GFP). **C)** Presses per sequence increased with training (main effect of training day: F_(6,192)_ = 89.34, p < 0.001), but differently across genotypes (main effect of genotype: F_(2,32)_ = 4.50, p < 0.05). Posthoc comparisons between genotypes confirm that presses/sequence were lower in KO-GFP mice compared to Con-GFP (p < 0.05) and normalized to controls in KO-D2 rescue mice (p = 0.99). **D)** The press rate increased throughout training (main effect of training day: F_(6,192)_ = 8.08, p < 0.001), but was not significantly different between genotypes in this experiment (F_(2,32)_ = 1.33, p = 0.28). **E)** Efficiency increased with training (main effect of training day: F_(6,192)_ = 14.46 p < 0.001) and differently between genotypes (F_(2,32)_ = 7.30, p < 0.01). Posthoc comparisons between genotypes demonstrate that KO-GFP animals had lower efficiency compared to controls (p < 0.01), and that was normalized in KO-D2 mice (p = 0.74). **F)** Sucrose reinforcers earned increased throughout training (F_(6,192)_ = 10.88, p < 0.001), differed between genotypes (F_(2,32)_ = 6.22, p < 0.01). Posthoc comparisons confirm that KO-GFP mice had lower levels of efficiency (p < 0.01) compared to Con-GFP, and that was normalized in KO-D2 mice (p = 0.88). **G)** There was a main effect of genotype for days to criteria (DTC; F_(2,32)_ = 10.70, p < 0.001). Posthoc comparisons demonstrate that KO-GFP mice had higher DTC compared to controls (p < 0.001), and that was normalized in KO-D2 (p = 0.93). **H)** In a progressive ratio task, the knockout groups performed differently from controls (F_(2,32)_ = 7.71, p < 0.001; one-way ANOVA). KO-GFP mice had a lower breakpoint compared to controls (p < 0.01, posthoc Tukey’s multiple comparisons test), which did not normalize to control levels in the KO-D2 group (p < 0.05 compared to Con-GFP, p = 0.59 compared to KO-GFP).

We hypothesized that the loss of D2Rs in ChIs of the Nucleus Accumbens (NAc), the ventral portion of the striatum attributed to reward learning, may be driving the deficit in progressive ratio performance. We injected the D2 rescue virus into the NAc of D2Flox-ChATCre mice and ran them through the progressive ratio task. We found that replacing D2Rs in NAc ChIs improved but did not completely rescue this behavior (Fig. S7).

## Discussion

We found that removing D2 receptors, and therefore removing a dopaminergic modulation mechanism in striatal cholinergic interneurons, has a profound effect on the learning, performance, and encoding of action sequences, without disrupting general locomotion or motor skill learning. Virally replacing D2 receptors in dorsal striatal ChIs rescued the sequence behavior, suggesting that dopamine modulation of ChIs is necessary for the learning, performance and encoding of action sequences. This D2 role could involve changes in transient ACh modulation of neuronal excitability and/or transmission, as well as altered synaptic plasticity.

There are numerous mechanisms by which ACh can influence MSN activity and behaviors involving the striatum. Activation of muscarinic receptors on MSNs can regulate their excitability. Both direct (D1-containing) and indirect (D2-containing) pathway MSNs express the muscarinic G_q_ coupled receptor M1, which enhances their excitability (*39–41*), while direct pathway MSNs preferentially express M4, an inhibitory G_i/o_ coupled receptor (*42, 43*). M1 receptors also contribute to dendritic excitability in indirect, but not direct pathway MSNs (*44*). The differential expression and activation of receptors suggests that ACh may modulate the two classes of MSNs in opposing directions, with direct pathway MSNs more likely to be inhibited, and indirect pathway MSNs more likely to be excited by acetylcholine. Likewise, optogenetically induced pausing of ChIs has been shown to both increase and decrease spontaneous firing of MSNs *in vivo* (*45, 46*). ACh can also indirectly inhibit MSNs through activation of nicotinic receptors on GABAergic interneurons (*47, 48*). Furthermore, both cortical and thalamic glutamatergic inputs to striatum contain M2 and M4 receptors that dampen glutamate release. Application of muscarinic agonists reduces glutamate release onto MSNs (*49, 50*), and can alter DA release (*51, 52*). Interestingly, driving a single action potential in a ChI can decrease EPSCs in neighboring MSNs, and application of muscarinic antagonists slightly increases glutamatergic transmission, indicating that ChIs provide a tonic inhibition of glutamatergic transmission in the striatum (*53*). Additionally, ACh activates nicotinic receptors on dopamine terminals to stimulate dopamine and co-neurotransmitter release (*54, 55*), and dopamine is an important regulator of multiple forms of striatal plasticity (*56–58*). Collectively these data indicate that the net effect of tonic acetylcholine is to dampen striatal transmission and support the idea that a brief pause may be necessary to disinhibit the circuit for plasticity and learning to occur. In support of this idea, artificially driving a pause in ChIs in the ventral striatum was shown to enhance associative learning in a fear conditioning paradigm (*59*). Overexpression of D2Rs in ChIs delayed acquisition of the NoGo rule in a Go/NoGo task, potentially due to larger cue-induced pauses in ChI firing that alter MSN activity resulting in impaired suppression of the Go response (27) (*29*).

Both ACh and D2 receptors have important roles in endocannabinoid-dependent long-term depression (eCB-LTD) at corticostriatal synapses (*60*). Activation of nicotinic receptors drives dopamine release that is necessary for eCB-LTD induction (*61, 62*), and LTD is disrupted in D2flox-ChATCre mice (*24*). This form of LTD is thought to be important for skill and instrumental learning, and aberrant plasticity is thought to play a role in the pathogenesis of movement disorders such as Parkinson’s and Huntington’s diseases (*56, 60*). While we cannot say that this specific form of LTD is involved in the operant sequence task used in this study, our lab has previously shown that CB1 receptors on cortical afferents are necessary for habitual action control using another operant paradigm (*63*). Specifically, eCB-CB1 signaling at orbitofrontal-dorsomedial striatum synapses is necessary to allow for action control by the dorsolateral portion of the striatum, the subregion implicated in control of habitual actions (*64, 65*). Thus, re-expression of Chi-D2Rs in DMS might contribute to rescue of action chunking in DLS. In the present study we did not specifically test if actions were habitual, but our sequence paradigm required extensive training, which is commonly used to promote habitual actions in rodents. The idea that dorsolateral striatum controls action sequencing in our task is also supported by our *in vivo* electrophysiological data. We found “chunking” MSNs, or those that changed their firing frequency throughout the entire sequence of lever pressing, only in the lateral portion of the striatum.

Dopaminergic cell firing precedes action initiation in mice performing a sequence of lever presses (*66*), and ChIs pause their firing during the action sequence (*3*). These studies suggest that the timing of dopamine release is primed to drive the pause in ChI firing and relieve the ACh-mediated tonic inhibition on the circuit, allowing for information transfer through and plasticity of striatal circuitry. However, other mechanisms for the pause in ChI firing have been suggested to have greater *in vivo* influence than the D2 action (*26, 46*), at least in anesthetized or head-fixed animals. Thus, it will be important to determine which mechanisms predominate during sequence behavior. Indeed, one of our goals was to determine if ChI physiology *in vivo* was changed in D2flox-ChATCre mice, but due the scarceness of ChIs, we were not able to do this using *in vivo* electrophysiology. We only detected 2 putative ChIs in recordings from 14 mice (Fig. S6). The difficulty of detecting ChIs in awake freely moving mice is evident by the lack of this data in the literature and a recent paper describing the difficulty of detecting ChIs using microendoscopes (*67*). As techniques improve, we will need to do further studies to examine the physiology of ChIs in relation to behavior. Indeed a large sample of putative ChIs that showed pauses following environmental stimuli was identified in a new study using advanced single-unit recording technology (*20*). Additionally, striatal ACh levels measured using a genetically encoded sensor showed transient decreases in ACh levels after reward predicting cues, and this decrease in ACh levels was reduced in D2flox-ChATCre mice (26) (*28*). In ventrolateral striatum, transient decreases in ACh were associated with certain phases of a reward-based decision making task that coincided with dopamine increases, and the ACh decreases were abolished in D2flox-ChATCre mice (30). In another study, calcium dynamics were measured with fiber photometry or imaging of soma and promixal dendrites of dStr ChIs expressing GCamp in head fixed mice provided evidence that ChIs show coordinated bursts of activity at transitions between movement states, as well as synchronization that decreased during sustained locomotion (*68*). In general, activity of DA afferent and ChI-burst calcium signals were strongly correlated at movement onset, but there was also a clear dissociation in that DA afferent signals persisted during continuous movement. It is not clear whether intracellular calcium measurements can detect brief pauses in ChI firing, as the authors of this paper acknowledge. However, decreases were apparent at the termination of spontaneous movement epochs and with deceleration. A burst-pause pattern was also observed following unexpected reward delivery, but was this was also associated with movement. Clear decreases were also observed around the initiation of DA increases when movements were not initiated. These findings suggest that ChI pauses related to DA signaling may contribute to the timing of action patterns, perhaps related to the pauses that have been associated with action cessation and orienting in primates (*69, 70*). While we were able to demonstrate loss of the DA-mediated pause and loss of plasticity in *ex vivo* slices in D2F-ChATCre mice (*24*), we can only speculate that we are blocking the DA-mediated pause and preventing learning associated plasticity in the striatum *in vivo*. Our knockout and rescue behavioral data indicate that D2-mediated regulation of dStr ChIs is important for learning action sequences, but the mechanisms are still unclear.

It should also be noted that a recent studies indicate that D2-mediated pauses are longer lasting or more readily detected in ChIs in slices from DMS compared to DLS, due to stronger glutamate release in DLS that produces bursts interrupting the D2-mediated pause (*71, 72*). While it is unclear if this is the case *in vivo*, this difference could factor into how D2 receptors influence the function of dStr regions implicated in goal-directed versus habitual actions. Activation of D2Rs may also inhibit ACh release by actions on ChI terminals (*50, 73*), and this mechanism could also contribute to the behavioral changes we observed.

A major strength of our data is that we were able to replicate the behavioral deficits in two independent D2flox-ChATCre lines. While problems with both of the ChAT-Cre lines have been reported (*32, 35*), we found that ChI physiology and behavior were not disrupted in these lines (*24*). We were careful to use only hemizygous Cre mice, thus minimizing the effect of transgene overexpression as much as possible. The similarity of our findings in both Cre lines and the rescue by viral D2 expression strongly indicate that the behavioral and neurophysiological alterations were due to loss of D2 function and not the Cre lines themselves.

The observation that D2 knockout in ChIs did not alter discrimination or reversal learning in the two lever task was somewhat surprising given previous findings suggesting that altering ChI numbers or function affects behavioral flexibility (*35–37*). These neurons have been implicated in switching of habitual responses (*74*). Certainly, loss of ChIs due to toxin-based lesions will produce much larger deficits in ACh signaling. Thus, it is not too surprising that this manipulation has effects not observed with D2 receptor loss. While our findings suggest that the D2Rs on ChIs are not crucial for flexibility in a reversal task, it may also be the case that our task was too simple to parse out more subtle differences in reversal learning, as mice were performing at chance levels on the first day of reversal.

The loss of ChI D2Rs did impair progressive ratio task performance. This may indicate a decrease in motivation or effort as task demand gets stronger. Alternatively, the knockout may promote rapid extinction of responding. The observation that the progressive ratio deficit was not rescued by D2 re-expression in either DS or NAc suggests that we did not target the correct brain area or combination of areas when trying to rescue this behavior with D2 re-expression. Perhaps re-expression is needed in all striatal subregions, or another brain region altogether. It should be noted that overexpression of D2Rs in NAc ChIs did not alter progressive ratio performance (*75*). It is also possible that this behavioral deficit was brought about by a developmental defect in neuronal activity or synaptic plasticity due to loss of D2 signaling in ChIs throughout the lifespan. These possibilities should be examined in future studies.

Our findings indicate an important role for D2 receptors on ChIs in the learning and performance of instrumental actions that becomes more prominent as the action requires greater effort. The loss of this modulatory dopamine action could contribute to problems in action performance in Parkinson’s disease, dystonia, antipsychotic-induced catalepsy and other movement disorders (*12, 13, 25, 76, 77*), as well as responses to drugs of abuse (*27, 78*). Thus, our observations may be useful in understanding, and ultimately treating, these disorders.

## Methods

### Animals

All experiments were approved by the Animal Care and Use Committee of the National Institute on Alcohol Abuse and Alcoholism and conformed to the NIH guidelines for care and use of experimental animals. Transgenic mice were bred in house. Mice were housed in groups of 2-4, except for mice with electrode implants, which were single housed, with ad libitum food and water unless otherwise stated. Animals were kept on a 12 hour light/dark cycle, with all experiments being performed during the light portion of the cycle. Both males and females were used. The mouse lines used were: D2flox (B6.129S4(FVB)-*Drd2^tm1.1Mrub^*/J, Jax #020631, (*79*); ChAT-IRES-Cre (B6N.129S6(B6)-*Chat^tm2(cre)Lowl^*/J, Jax #018957); Tg-BAC-ChATCre (*80*); TdTomato reporter (B6.Cg-Gt(ROSA)26Sortm14(CAG-TdTomato)Hze/J; Jax #007914).

### Adapted self-paced FR8 instrumental lever-pressing paradigm

Adult (8-20 week old) mice were food restricted to 85-90% of their body weight prior to training by feeding them ∼10% of their body weight for 2-3 days. This low body weight was maintained early in training, while later in training (FR8 stage) body weight was maintained between 90-95% of the starting weight. Mice were placed in operant chambers housed in sound-attenuating boxes (Med-Associates, St. Albans, VT). The first day of training consisted of a 30 min pretraining session, in which no lever was present, and sucrose (30-50 µL, 20%) was delivered via syringe pump into a cup receptacle on a random interval schedule, with one delivery per 60 seconds on average (RI60). For the reminder of the training sessions, the house light was illuminated and a single lever (retractable ultrasensitive mouse lever) extended at the onset of the session that remained extended with the house light on throughout training. The first four days of operant training were on a fixed ratio (FR) 1 schedule, in which a single lever press was reinforced with sucrose. The maximum number of reinforcers increased with the FR1 training session, with 10 allowed on day 1, followed by 15 on day 2, and 30 on days 3-4. All FR sessions were 60 minutes maximum, and animals remained at each training level until they reached a criterion of 15 reinforcers/60 minute session. Animals that received at least 15 reinforcers on day 4 of FR1 were moved to continuous FR4 training, in which the 4^th^ lever press of a pressing sequence was reinforced, where a sequence was defined as continuous presses without intervening licking, measured using contact lickometers. Mice remained at the FR4 stage for a minimum of 4 days but remained at this stage until reaching criterion. Mice were then moved to the continuous FR8 stage, in which the 8^th^ press in a sequence was reinforced. Mice were trained at the FR8 stage for 8 days or more until reaching the criterion of 15 reinforcers in a one-hour session. FR4 and FR8 sessions had no maximum number of reinforcers.

### Progressive ratio task

A self-paced progressive ratio (PR) task, similar to that described in Richardson and Roberts, 1996 (*81*), was performed following the completion of the sequence training. Mice were trained on a standard FR10 schedule until they received at least 15 sucrose rewards within an hour on 2 consecutive days. On the following day, the number of lever presses needed to earn a reward increased, until the number of lever presses needed to achieve a reward was not reached within an hour. The breakpoint is defined as the next reward requirement the mouse failed to achieve.

### Reversal task

Mice were food restricted to 85-90% of their body weight, then trained in operant boxes with two levers present, in which each lever was reinforced with ∼30 µL 20% sucrose, for a minimum of 4 days or until they reached at least 30 reinforcers in a one-hour session for two consecutive days. Then they moved into the acquisition phase, in which only one lever was reinforced in a biased design. All mice preferred one lever (> 60% of presses on that lever). The nonpreferred lever from the training phase became the reinforced lever in the acquisition phase, while the preferred lever was not reinforced. After 8 days of acquisition, the levers were reversed, and mice were trained for 8 days with the opposite lever being reinforced on an FR1 schedule.

### Accelerating rotarod

Mice were initially habituated to the behavior room for 1 hour the day before training. Animals were placed on a rotating rod (5 lane Rota-rod for mouse, Med Associates), which gradually accelerated from 4 to 40 rpm over a five min trial. The trial ended when the mouse fell of the rod, which was detected by beam breaks and collected using Rota-rod software (Med Associates), or at the end of 5 minutes. The experimenter manually stopped the trial if the animal began to ride around the rod. Animals underwent three trials per day with a 20 min inter-trial interval (ITI) across four days of testing.

### Locomotor activity

Mice were placed in a novel clear polycarbonate chamber (20 cm H x 17 cm W x 28 cm D), which was placed in an infrared photobeam detector (Columbus Instruments, Columbus, OH). Locomotor activity was measured as beam breaks per 10 seconds for a one-hour session.

### In vivo extracellular recordings

Mice were anesthetized with isoflurane and implanted with 16 channel tungsten multi-electrode arrays (50 µm diameter of individual electrodes; 8 electrodes in 2 rows; 200 µm electrode spacing; 1250 µm row spacing; Innovative Neurophysiology, Durham, NC) in the dorsal striatum, such that one row targeted the dorsal medial, and the other targeted the dorsal lateral striatum (Figure 5A). The center point of the array from bregma was 0.5 mm rostral and 1.75 mm lateral, and the electrodes were lowered 2.25-2.5 mm from the brain surface. The electrodes were grounded with a silver wire to 2 stainless steel screws implanted supradurally in the skull and secured using cold-cure dental cement. Mice were given 5mg/kg ketoprofen i.p. and fluids 2 days post-surgery and allowed to recover 2-4 weeks.

Prior to behavioral testing, mice were given 2-3 habituation sessions, in which they were briefly anesthetized with isoflurane and connected to a 12-channel headstage (20x gain; Plexon Inc., Dallas, TX), then placed in their home cage while units were initially sorted using Sort Client, an online sorting algorithm (Plexon, Inc.). During behavioral testing, mice were again briefly anesthetized to be connected to the headstage, and were allowed to recover one hour prior to training, in which units were presorted using the online sorter. During training, unit activity was collected using the MAP system (Plexon, Inc.) at a digitization rate of 40 kHz with 12-bit resolution with a 250 Hz highpass filter. TTL pulses from the Med-Associates system were sent to the MAP system through an A/D board (Texas Instruments Inc., Dallas, TX) to timestamp behavioral events with unit recordings. Data were then resorted with Offline Sorter (Plexon, Inc.) using principal component analysis to identify single units determined by multivariate ANOVA in 3D cluster space and pairwise 3D single unit comparisons. Units were classified using baseline firing rate, peak to trough amplitude and half-width (Fig. S6, and activity of putative MSNs was aligned to behavior.

To align units to behavior, we constructed peri-event histograms in NeuroExplorer (Nex Technologies, Madison, AL) around the time-stamped event (lever press, first lever press, etc.), in which neural activity was averaged in 20 ms bins distributed from 2 seconds before to 2 seconds after the event, with 95% confidence intervals. Units were considered task related if the firing rate increased by 95% or decreased to 95% of baseline within 500 ms of the event. Units were considered start/stop if they were found to be task responsive to first and/or last press, but not intervening presses, whereas “chunking cells” met the criteria for changes in firing throughout the entire duration of the sequence. Units were considered press responsive if they were found to be task responsive to the lever press but were not considered start/stop or sustained.

### Stereotaxic viral injections

We injected AAV2/1-hSyn-DIO-D2R(L)-IRES-mVenus (*38*) or the control virus, AAV2-hSyn-DIO-eGFP (UNC Vector Core, Chapel Hill, NC), bilaterally into the dorsal or ventral striatum of D2flox or D2flox-ChATCre littermates. Dorsal coordinates: AP, +0.6, ML, +/ − 2.5; DV −3.0 and −2.2. Ventral coordinates: AP, +1.7 mm; ML, ±1.2 mm; DV, −3.9 mm. 200 nl of virus was injected at each site at a rate of 50 nl/min. Behavioral experiments were run 5-8 weeks post-injection.

### Histology

Mice were transcardially perfused with PBS then 4% formaldehyde. Brains were removed and post-fixed in 4% formaldehyde overnight, then sliced with a Pelco easiSlicer vibratome (Ted Pella Inc., Redding, CA) into 50 μm slices. Slices were subjected to the ‘Brain BLAQ’ protocol (*82*) before staining, then blocked in PBST with 5% BSA overnight. For electrode implanted animals, slices were stained with GS-IB_4_-Alexa Fluor 488 Conjugate (1 μg/mL; Invitrogen, Carlsbad, CA #I21411), propidium iodide (1 μg/mL; Molecular Probes, Eugene, OR), and DAPI (0.25 μg/mL; Invitrogen) in blocking solution overnight (Figure 4A). For viral injected animals, the virally expressed GFP or YFP was enhanced using the rabbit anti-GFP Tag Polyclonal Antibody-Alexa Fluor 488 (Invitrogen #A21311). In the viral rescue experiment (Figure 5A-B), slices were also stained with a goat anti-ChAT primary antibody (1:500; #AB144P, Millipore, Burlington, MA), and Donkey anti-Goat secondary antibody-Alexa Fluor 594 (1:1000, #A-11058, Invitrogen). Slices were washed in PBST four times over a 16-24 hour period, then 3 times over a one hour period in PBS. Slices were mounted in Fluoromount (Sigma-Aldrich, St. Louis, MO) and imaged using an AxioZoom.V10 microscope (Zeiss).

### Data Analysis

Statisical analyses were done using Prism (GraphPad Software, Inc., La Jolla, CA) or STATISTICA 12 (Dell Software, Redmond, WA). Data are presented as means +/- SEM. The specific tests performed are described in the results and/or figure legend for each experiment. Differences were considered significant when p < 0.05.

## Acknowledgements

We would like to thank Dr. Karina Abrahao for assisting with viral rescue surgeries and other members of the Laboratory for Integrative Neuroscience for helpful comments and discussions. This work was funded by the NIAAA Division of Intramural Clinical and Biological Research (ZIA AA000416) and R01MH54137 to JAJ that funded the production of the D2 rescue virus.

## Footnotes

Author contributions: J.H.C. and D.M.L. designed the study and wrote the paper. J.H.C performed and analyzed all experiments. C.K. and J.A.J. provided the D2 virus, helped design viral rescue experiments and edited the paper.

