## Supplemental Figures 1-7 for "Dopaminergic D2 receptor modulation of striatal cholinergic interneurons contributes to sequence learning"

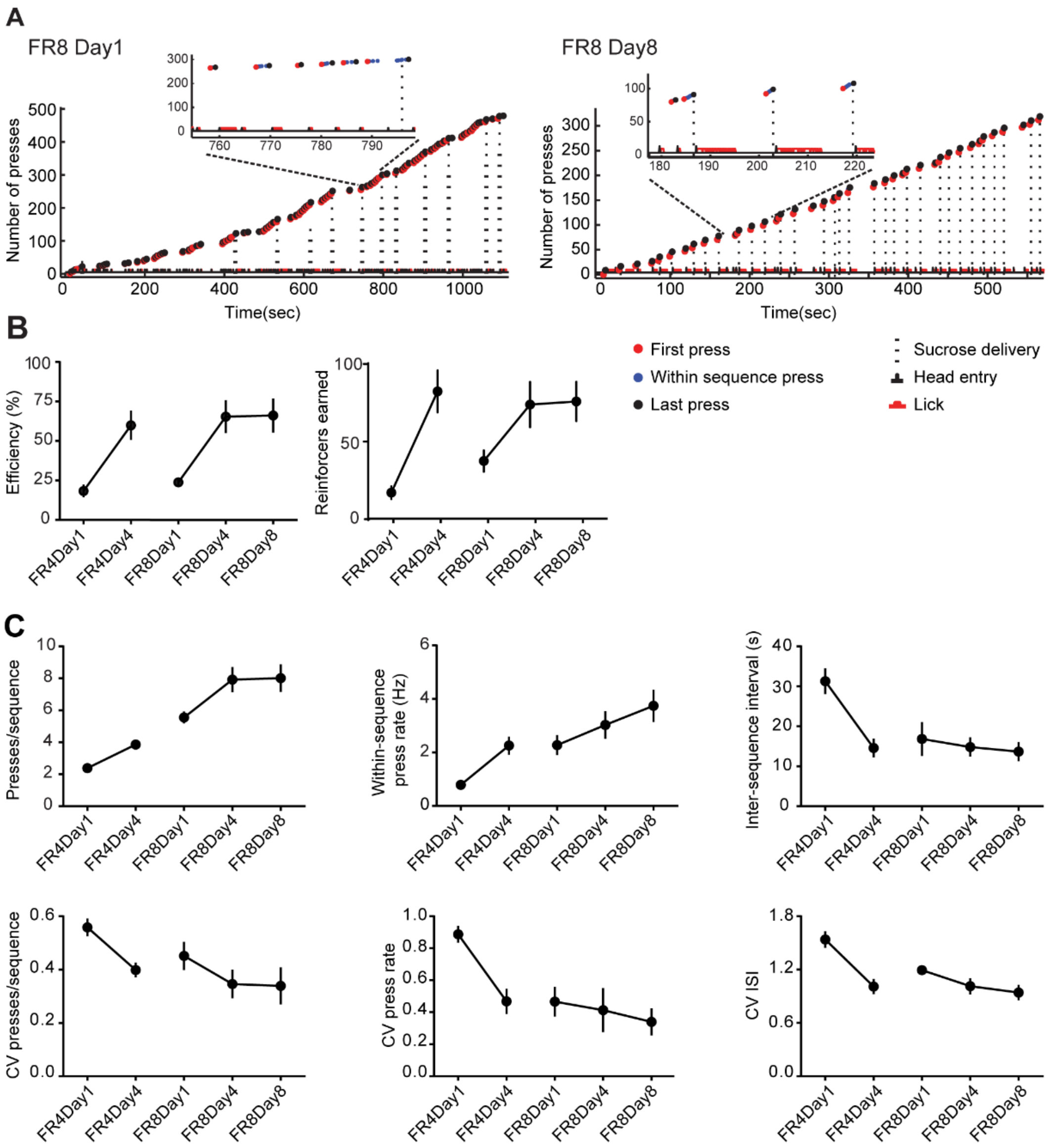


**Supplementary Figure 1. Modified FR8 sequence task promotes consistent action sequences in C57/B6 mice. A)** Cumulative lever press plots showing examples of behavioral shaping from days 1 and 8 of FR8 training in C57/B6 mice. **B)** Efficiency (# of sequences/reward *100; $F_{(4,35)} = 8.39$, $p < 0.001$, one-way ANOVA) and number of reinforcers earned ($F_{(4,35)} = 6.58$, $p < 0.001$) increased with days of training. **C)** Learning and consistent action sequences are demonstrated by increases in presses/sequence

($F_{(4,35)}$ = 18.99, $p < 0.001$) and the within-sequence press rate ($F_{(4,35)}$ = 6.70, $p < 0.001$), and a decrease in the inter-sequence interval ($F_{(4,35)}$ = 6.01, $p < 0.001$), as well as a decreases in the within mouse coefficient of variation of those measures ($F_{(4,35)} > 3.31$, $p < 0.05$) across training days. N = 8 mice.

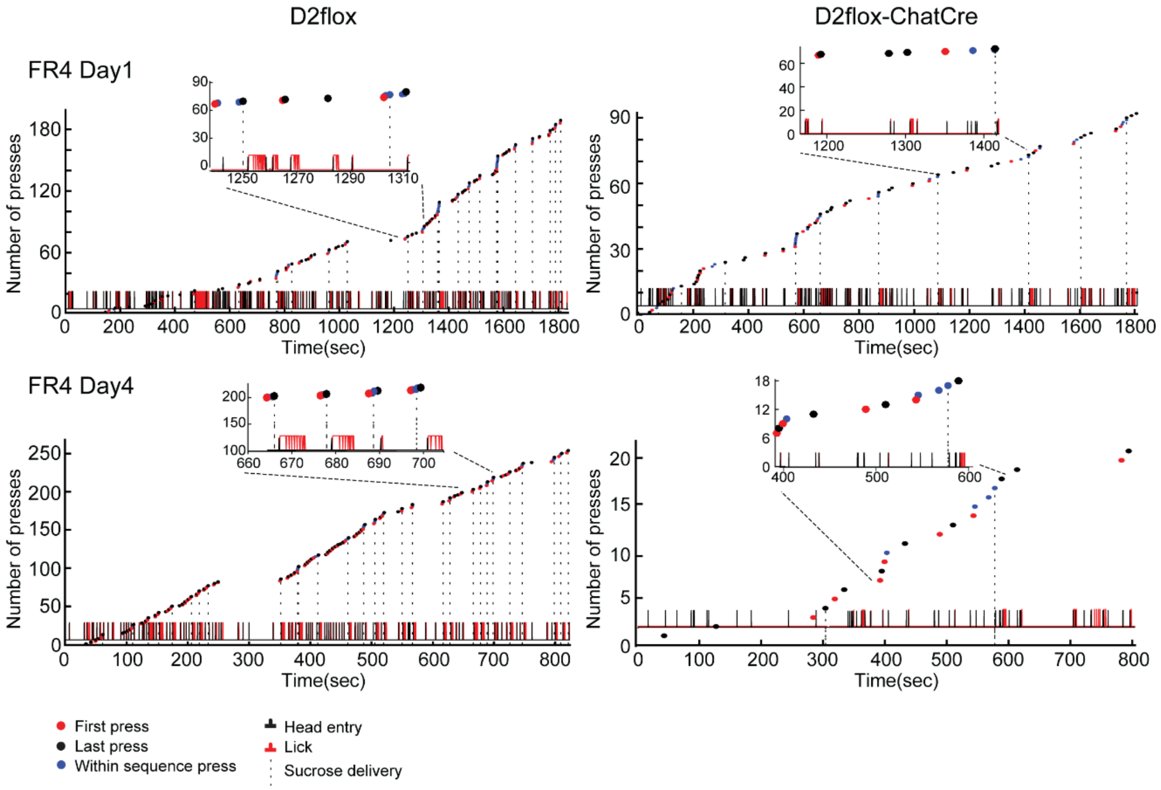


**Supplementary Figure 2. Examples of lever pressing in the FR4 portion of the sequence learning task.** Cumulative lever press plots showing examples of behavioral shaping in D2flox (left) and D2flox-ChATCre (right) littermates performing the adapted FR task. Top: The first day of FR4 training. Bottom: 4$^{th}$ day of FR8 training in which the D2flox-ChATCre animal presses less frequently, slower, and with lower efficiency than the D2flox littermate.

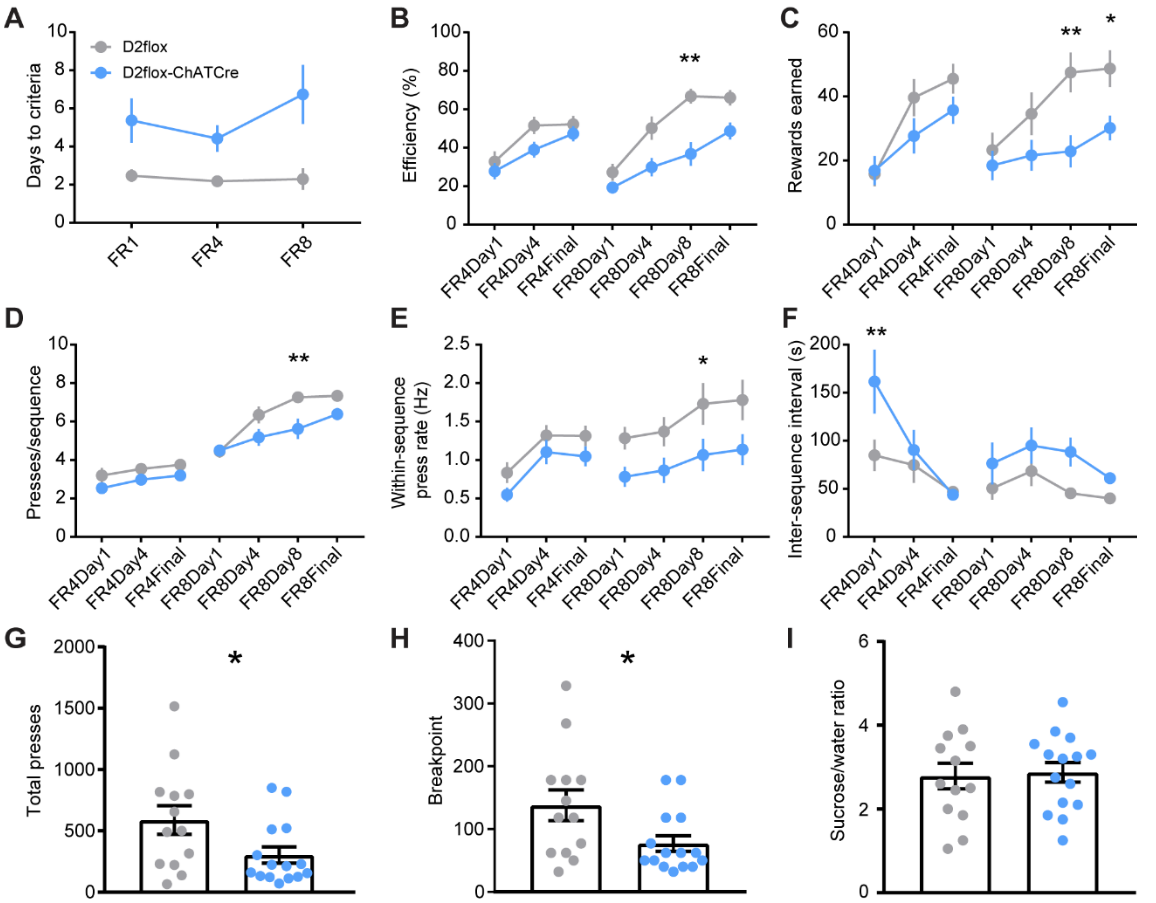


**Supplementary Figure 3. D2flox-BAC-Tg-ChATCre have deficits in sequence learning and progressive ratio task. A)** D2flox mice crossed with Tg-BAC-ChATCre mice required more training than D2flox littermates to reach the criterion at all levels of training (main effect of genotype: $F_{(1,34)} = 18.37$, $p < 0.001$). **B)** They were able to reach a level of efficiency similar to littermates, but at a different rate (main effect of day: $F_{(6,204)} = 25.35$, $p < 0.001$; genotype: $F_{(1,34)} = 9.03$, $p < 0.01$; interaction: $F_{(6,204)} = 36.00$, $p < 0.01$). **C)** They received fewer sucrose rewards than littermates (main effect of day: $F_{(6,204)} = 13.34$, $p < 0.001$; genotype: $F_{(1,34)} = 4.86$, $p < 0.05$; interaction: $F_{(6,204)} = 2.79$, $p <0.05$). **D)** The presses per sequence increased throughout training ($F_{(6,204)} = 79.83$, $p < 0.001$), but at a different rate in D2flox -ChATCre animals than D2flox littermates (main effect of genotype: $F_{(1,34)} = 6.38$, $p < 0.05$; interaction: $F_{(6,204)} = 2.15$, $p < 0.05$). **E)** The within sequence press rate was slower (main effect of genotype: $F_{(1,34)} = 4.37$, $p < 0.05$; main effect of day: $F_{(6,204)} = 13.39$, $p < 0.001$ ) and inter-sequence interval was longer (main effect of genotype: : $F_{(1,34)} = 5.32$, $p < 0.05$; main effect of day: $F_{(6,204)} = 5.82$, $p < 0.001$) in knockouts compared to littermates. **A-E)** We performed two-way repeated measures ANOVAs, and Sidak's multiple comparisons tests between genotypes at each stage of training (n = 17 D2flox and 19 D2flox-ChATCre). Significant differences in the multiple comparisons are indicated by asterisks (* p < 0.05, ** p < 0.01). **F)** In a progressive ratio task, D2flox-ChATCre mice pressed fewer times (p = 0.03) and had lower breakpoints (**G;** p = 0.03) than D2flox littermates. **H)** This reduction in breakpoint performance was not due to a reduced sucrose preference, because there was no difference in a home cage sucrose preference test between the knockouts and littermates (p = 0.82). **F-H)** Student's t-tests were performed. N = 13 D2flox and 15 D2flox-ChATCre.

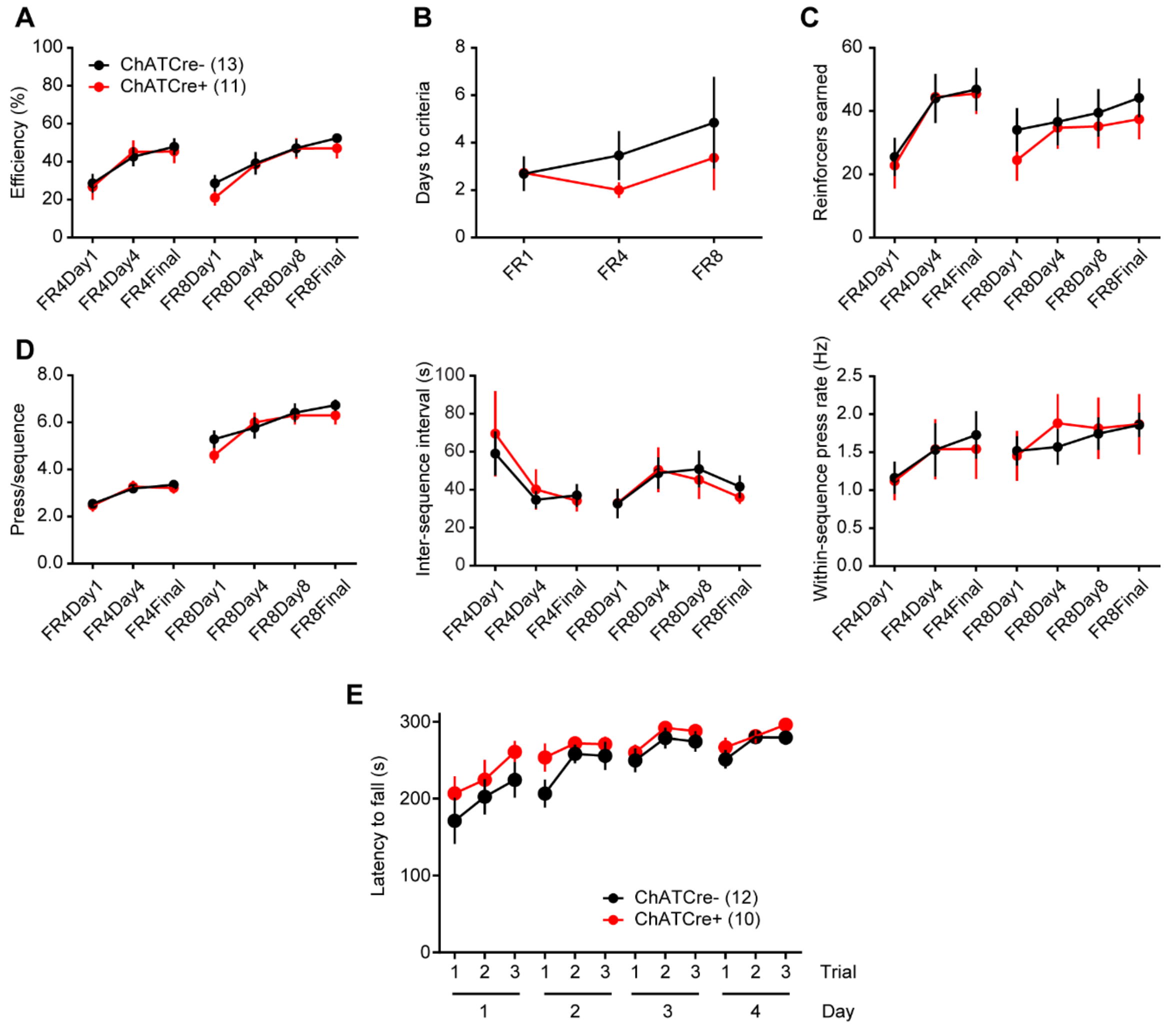


**Supplementary Figure 4. BAC-Tg-ChATCre mice performed similar to Cre- littermates in the FR8 sequence learning task. A)** ChATCre mice (red) reached the same levels of efficiency at all stages of training as Cre- littermates (black; $F_{(1, 22)} = 0.161$, $p = 0.69$), and both genotypes increased efficiency throughout training ($F_{(6, 132)} = 16.2$, $p < 0.0001$; n values in parentheses). **B)** ChATCre mice and littermates also required similar amounts of training to reach criteria ($F_{(1, 22)} = 0.938$, $p = 0.34$) and **C)** earned a similar number of reinforcers ($F_{(1, 22)} = 0.183$, $p = 0.67$), which increased with training ($F_{(6, 132)} = 10.8$, $p < 0.001$). **D)** The number of presses per sequence, inter-sequence interval, and press rate did not differ between genotypes ($F_{(1, 22)} < 0.233$, $p > 0.64$). **E)** ChATCre animals perform similar to Cre- littermates in an accelerating rotarod test. Both genotypes learned the task (main effect of trial: $F_{(11, 220)} = 10.74$, $p < 0.001$) with no effect of genotype ($F_{(1, 20)} = 2.56$, $p = 0.13$).

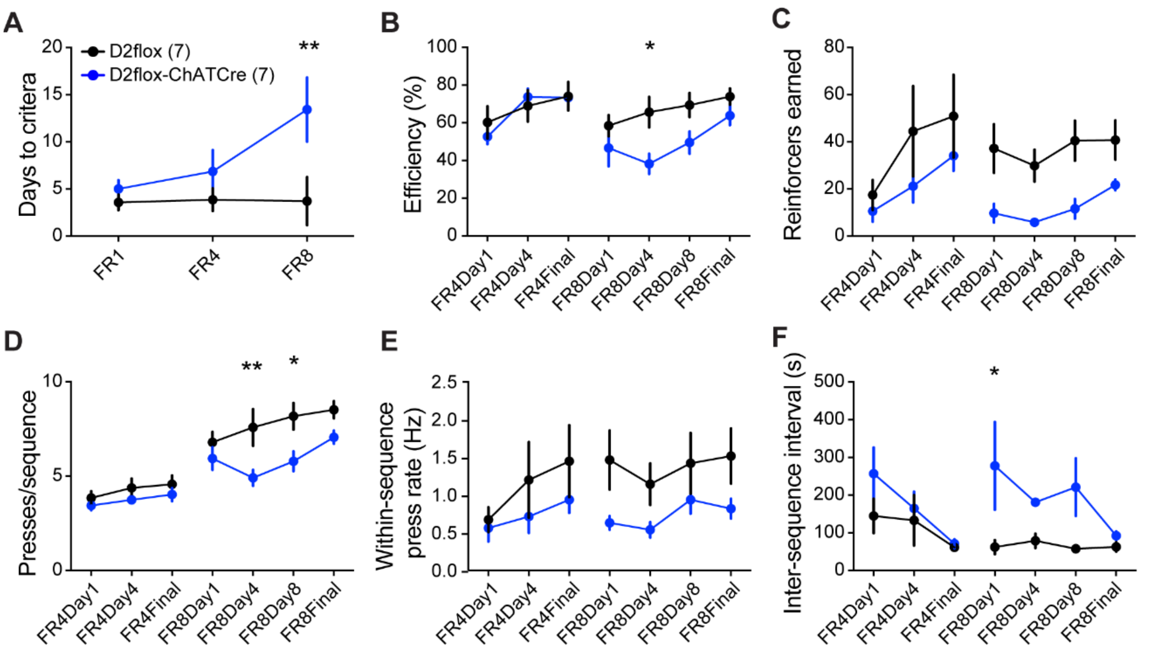


**Supplementary Figure 5. Implanted D2flox-ChATCre mice show deficits in sequence learning compared to implanted D2flox littermates. A)** Implanted D2flox -ChATCre (blue) mice required more training than implanted D2flox littermates (black) to reach the criterion (main effect of genotype: $F_{(1,12)} = 6.203$, $p < 0.05$). **B)** They were able to reach a similar level of efficiency as littermates, but at a different rate (main effect of day: $F_{(6,72)} = 6.363$, $p < 0.001$; genotype: $F_{(1,12)} = 2.568$, $p = 0.135$; interaction: $F_{(6,72)} = 2.274$, $p < 0.05$). **C)** They received fewer sucrose rewards than littermates throughout training (main effect of day: $F_{(6,72)} = 3.623$, $p < 0.01$; genotype: $F_{(1,12)} = 5.669$, $p < 0.05$; no significant interaction: $F_{(6,72)} = 0.552$, $p = 0.7$). **D)** The presses per sequence increased throughout training ($F_{(6,72)} = 32.94$, $p < 0.001$), but at a different rate in knockouts compared to controls (main effect of genotype: $F_{(1,12)} = 6.46$, $p < 0.05$; interaction: $F_{(6,72)} = 2.684$, $p < 0.05$). **E)** The within-sequence press rate was not significantly different between genotypes in implanted animals (genotype: $F_{(1,12)} = 2.248$, $p = 0.16$), but **(F)** the inter-sequence interval was longer in knockouts compared to controls (main effect of genotype: $F_{(1,12)} = 11.95$, $p < 0.01$) **A-F)** We performed two-way repeated measures ANOVAs, and Sidak's multiple comparisons tests between genotypes at each stage of training. N values are in parentheses. Significant differences in the multiple comparisons are indicated by asterisks (* $p < 0.05$, ** $p < 0.01$).

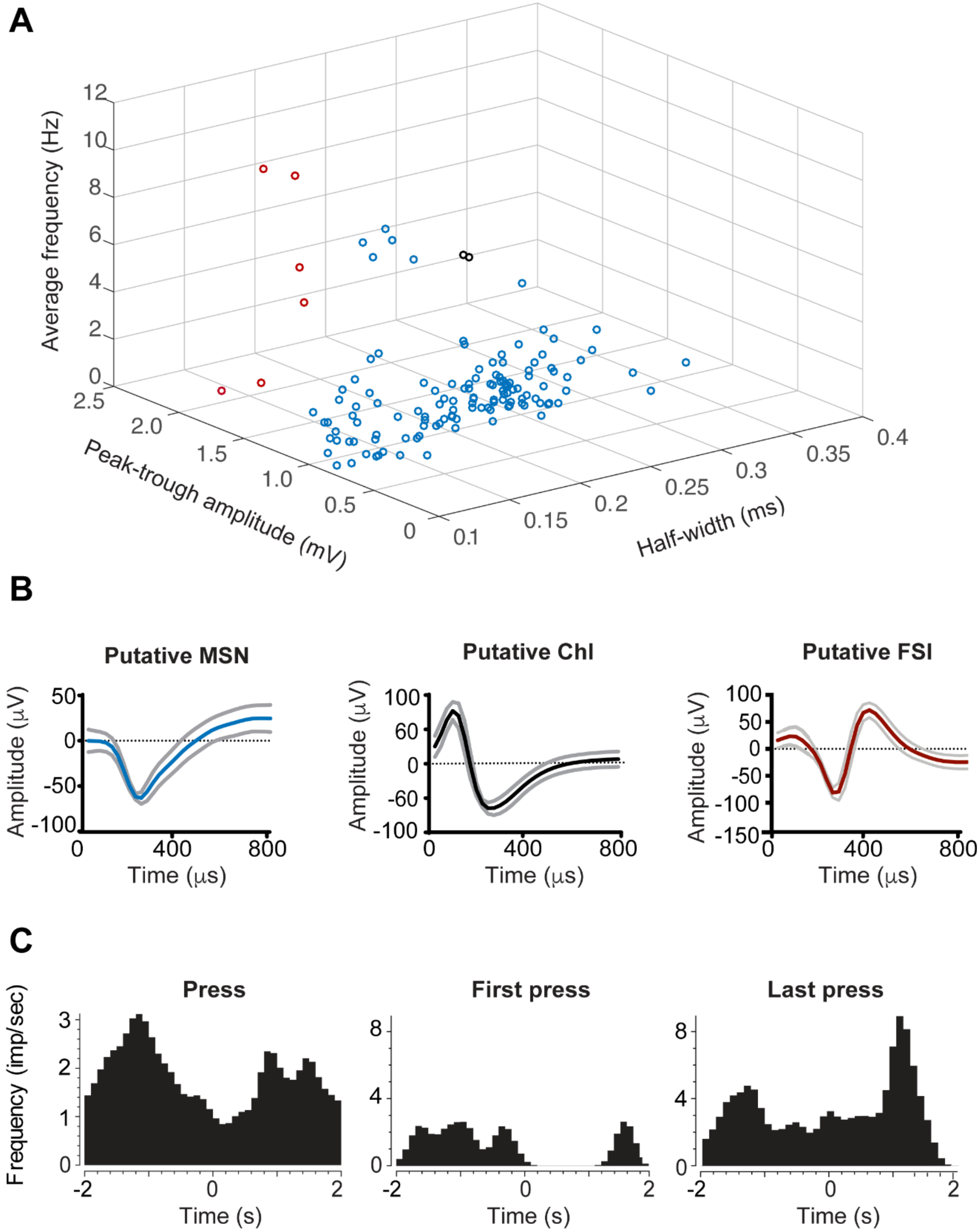


Supplementary Figure 6. Classification of units. A) Units were classified based on average firing frequency, peak to trough amplitude, and waveform half-width. Units with large amplitudes (> 2 mV) and short half-widths (< 0.2 ms) were classified as putative fastspiking interneurons (red). Units with half-widths > 0.25 ms and frequencies > 2 Hz were classified as putative ChIs (black). All other units were considered MSN (blue). **B)** Example waveforms displayed as averaged traces with SD in gray. **C)** Of the three putative ChIs, 2 were from D2flox and one was from a D2flox-ChATCre mouse. One of two WT ChIs was task responsive. The perievent histogram for that unit is shown here. It paused at the start of lever pressing and bursted after the termination of pressing.

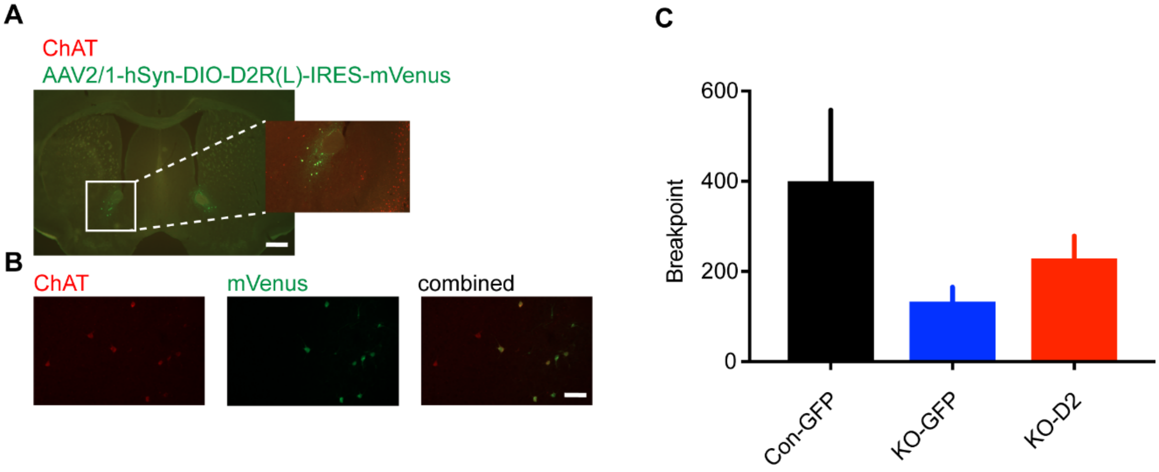


**Supplementary Figure 7. No difference between groups in the progressive ratio task when D2Rs are expressed in ventral striatal ChIs of D2Flox-ChATCre mice.** **A)** Example of AAV2/1-hSyn-DIO-D2R-IRES-mVenus virus (green) injected bilaterally into the ventral striatum of a D2flox-ChATCre mouse that was stained for choline transferase (ChAT, red). Scale = 500 μm. **B)** Viral expression was limited to $ChAT^{+}$ cells. Scale = 50 μm. **C)** The breakpoint values are 399 ± 159 for D2flox mice injected with the control virus (Con-GFP; black; n = 9), 132 ± 33 for D2flox-ChATCre injected with the control virus (KO-GFP; blue; n = 7), and 228 ± 51 for D2flox-ChATCre injected with the D2 virus (KO-D2; red; n = 9). The breakpoints are not statistically different across groups ($F_{(2,22)} = 1.59$, $p = 0.2$; one-way ANOVA).
